# News without the buzz: reading out weak theta rhythms in the hippocampus

**DOI:** 10.1101/2023.12.22.573160

**Authors:** Gautam Agarwal, Brian Lustig, Seiji Akera, Eva Pastalkova, Albert K. Lee, Friedrich T. Sommer

## Abstract

Local field potentials (LFPs) reflect the collective dynamics of neural populations, yet their exact relationship to neural codes remains unknown^1^. One notable exception is the theta rhythm of the rodent hippocampus, which seems to provide a reference clock to decode the animal’s position from spatiotemporal patterns of neuronal spiking^2^ or LFPs^3^. But when the animal stops, theta becomes irregular^4^, potentially indicating the breakdown of temporal coding by neural populations. Here we show that no such breakdown occurs, introducing an artificial neural network that can recover position-tuned rhythmic patterns (pThetas) without relying on the more prominent theta rhythm as a reference clock. pTheta and theta preferentially correlate with place cell and interneuron spiking, respectively. When rats forage in an open field, pTheta is jointly tuned to position and head orientation, a property not seen in individual place cells but expected to emerge from place cell sequences^5^. Our work demonstrates that weak and intermittent oscillations, as seen in many brain regions and species, can carry behavioral information commensurate with population spike codes.

---

Recording from an electrode in the brain reveals local field potentials (LFPs) that reflect the coordinated activity of many thousands of neurons. While oscillations in the LFP are a widespread phenomenon ^6,7^, their role in brain function remains debated ^8^. Proponents point to their computational potential ^9–11^, while others question if they are merely epiphenomenal ^12^.

The theta rhythm of the rat hippocampus is one of the most extensively studied oscillations. Within this region, neurons called place cells fire when a rat occupies specific locations of its environment, known as the cells’ place fields. When the animal traverses a place field, the cell fires rhythmic bursts at progressively earlier phases of the ongoing theta rhythm, a phenomenon known as phase precession ^2^. A consequence of phase precession is that, within a single theta cycle, place cells fire sequentially according to the order in which their place fields are traversed (see Fig.2 of ^13^). The theta rhythm can be viewed as a communication channel carrying information contained in the rhythmic spiking of place cell populations to other brain regions.

In previous work, we investigated whether position information could be derived from the theta rhythm of the LFP, reflecting the collective activity of place-tuned cells. We found anatomically distributed phase and amplitude patterns in the theta frequency band (5-11 Hz) that indicated the rat’s current location with an accuracy comparable to traditional spike-based decoders ^3^. However, our approach hinged on using the theta rhythm as a reference for reading out LFP phase patterns. This limits its generalizability, as the strength of theta varies across behavioral conditions ^4^ and species ^14–16^. Here, we investigate whether positional information can be recovered from theta rhythms when they are weak and irregular. One possibility is that a weak theta rhythm reflects a lack of coherence among place cells, resulting in noisy LFPs that do not carry positional information. Instead, we find that position information can be recovered from LFPs when theta is weak, but this is most effective when we detect place-selective patterns at theta frequency (‘place theta’ or ‘pθ’) without using the dominant, classically observed theta rhythm (‘θ’) as a reference frame. While a dominant brain rhythm might naturally appear to be a pacemaker organizing a neural code, it is not always best for reading out the code.

## Results

### Carrier-based decoding is less reliable during immobility

We first attempt to decode position information from LFPs under the assumption that position modulates a single theta rhythm, analogous to how information is carried by a radio signal. We recorded LFPs from dorsal hippocampal region CA1 using up to four 64-shank arrays while rats navigated a 3-arm maze to collect water rewards (Fig. 1 A-B). This task structure allowed us to sample periods when the theta rhythm is strong and consistent (i.e., during running, Fig 1C left), as well as when it is weak and inconsistent (i.e., during staying, Fig. 1C right; Fig. 1E). As proposed previously ^3^, we 1) transform the time series measured at each electrode into a complex-valued analytic signal with a defined phase and amplitude ^17^; 2) use PCA to identify a single dominant component ‘θ’ that serves as the carrier; 3) demodulate the LFP by subtracting the phase of θ from the local phase at each electrode, thereby removing the oscillatory component of the signal; and 4) train a linear classifier on the demodulated LFP to predict which maze arm is being occupied by the animal (Fig. 1D). During periods of strong θ, the demodulated LFP has a more consistent form that accurately predicts the animal’s position (Fig. 1C-D, left)^3^. By contrast, when the rat stops running, the demodulated LFP becomes less consistent and is less predictive of the animal’s location (Fig. 1C-F).

**Fig. 1.**
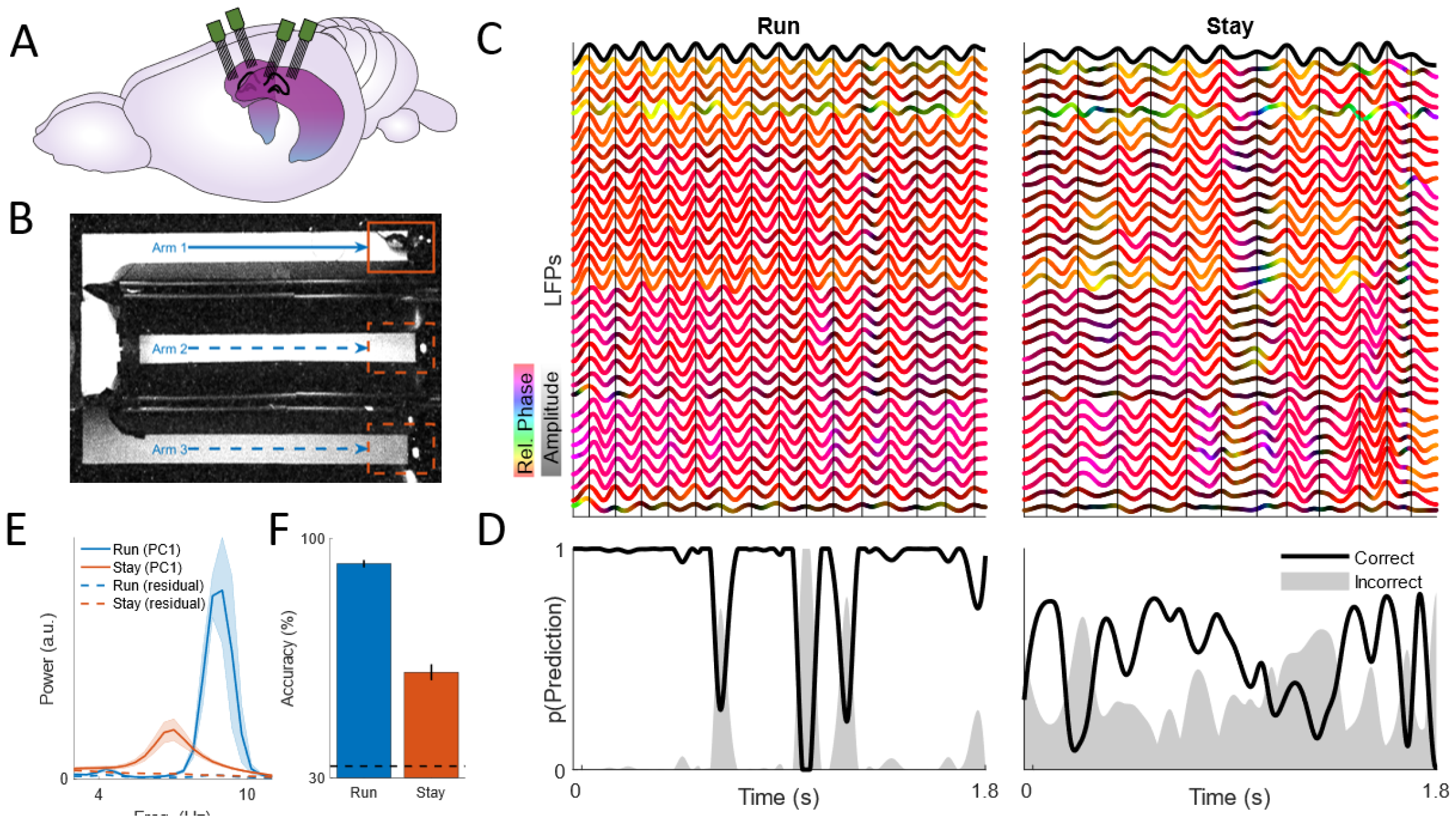
Carrier-based decoding during strong and weak theta. **A)** Neural activity was recorded from up to 4 64-shank multi-electrode arrays implanted bilaterally in dorsal CAI. Image downloaded and modified from scidraw.io. **B)** Rats were trained to sequentially visit ports in a 3-arm maze to receive water reward, allowing us to sample LFPs during running (blue) and staying (red). **C)** Theta-band LFPs recorded during running (left) and staying (right), respectively showing strong and weak theta rhythms. The top black line indicates ‘0’, the first principal component of the multi-channel LFP. Subsequent lines correspond to LFPs measured at individual electrodes and color-coded based on their instantaneous phase relative to θ (yellow - leading, purple - lagging) and amplitude (black - low, colored - high). Vertical lines indicate θ peaks. LFPs show greater amplitude and more consistent phase relationships during running than during staying. **D)** A linear classifier predicts arm occupancy during running more consistently than during staying. **E)** θ power spectra during running (blue) and staying (red) across over 4 sessions. **F)** Carrier-based decoding is more accurate during running than staying across sessions. **E** and **F** depict mean ± s.e.m., n = 4 sessions from 3 rats.

### Carrier-based decoding is susceptible to shared noise

To examine why the decoding precision is reduced during staying, we tested a simple model of theta LFP generation (Extended Data Fig. 2). The model consists of a large population of place cells whose place fields tile the environment uniformly (Fig. 2A, top). These cells are phase-locked to a theta carrier in a position-dependent manner ^2,18,19^. The LFP at different electrodes is modeled as a linear superposition of neural responses (Fig. 2B, top). The resulting simulated LFPs exhibited slight place-dependent modulations in amplitude and phase that are due to inhomogeneities in electrodes’ sampling of place cell activity and that resemble those found experimentally (Fig. 1C). We can then use linear regression to derive a collection of decoders that detect the multielectrode pattern unique to each position along the track (‘pθ’, Fig. 2C, top). We can estimate the animal’s position as the one corresponding to the most active pθ. Because the decoders produce complex-valued outputs, pθ activation can be read out in different ways. One approach is “carrier-based decoding”, in which the real component of the demodulated pθ activations is compared (Fig. 2D, top). This approach is sensitive to the phase of each pθ relative to the theta carrier. A second approach is “carrier-free decoding”, in which the magnitude of pθs is compared, without any demodulation (Fig. 2E, top). For the noiseless LFP model, carrier-based decoding is superior to carrier-free because the LFPs’ phase relative to the theta carrier contains information about position (Fig. 2C-F, top). Note that under noiseless conditions, the two methods perform equally well if sufficiently many electrodes are used.

**Fig. 2.**
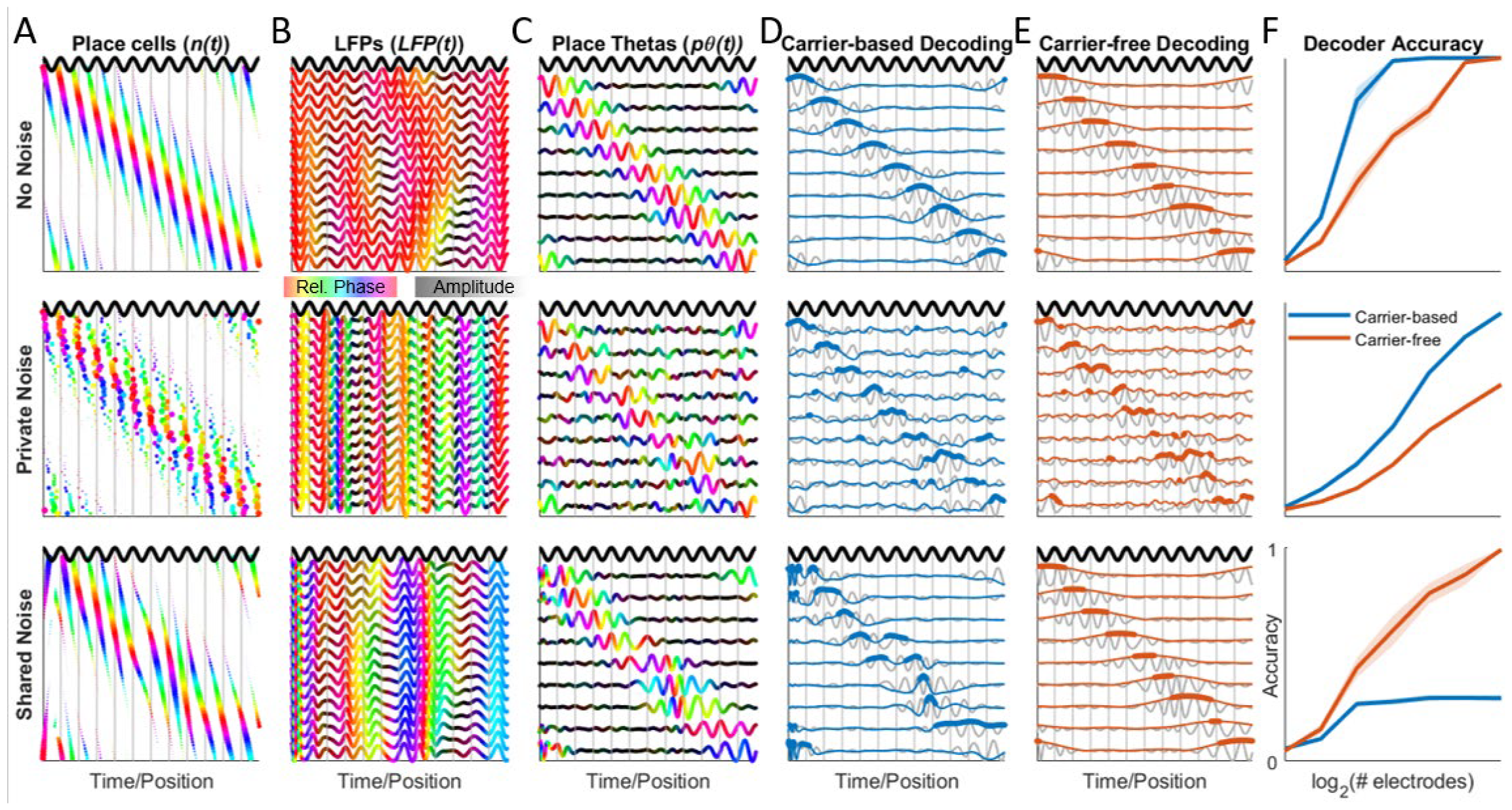
LFP decoding strategy depends on type of noise. *Top row*: responses of model neurons contain no noise. *Middle row*: the phase of each neuron is perturbed independently (‘private noise’). *Bottom row* - the phase of all neurons is perturbed together (‘shared noise’). **A)** Place cell responses for a single traversal of a linear environment. Dot size indicates magnitude of response while color indicates phase of response relative to theta (dark black oscillation). Vertical gray lines indicate theta peaks. **B)** Individual electrode responses (LFP(t)) during the same trajectory. **C)** Position-varying LFP patterns in B can be linearly transformed to yield place-tuned theta (p0) oscillations. In **B-C)**, data is complex-valued, with color indicating relative phase and intensity representing amplitude (as in Fig. **1C**). **D)** Carrier-based decoding involves taking the real projection of the p0s following demodulation by θ (blue lines) and selecting the largest one at each time step (blue dots). **E)** Carrier-free decoding involves taking the absolute value of the p0s and selecting the largest one at each time step (red dots). **D-E)** Gray traces show p0s duplicated from **C. F)** *Top*: Carrier-based decoding is superior when using fewer electrodes because it is more spatially selective than carrier-free decoding (i.e., the blue lines in **D** are more peaked than the red lines in **E)**. *Middle*: With private noise, carrier-based decoding remains superior. *Bottom*: With shared noise, carrier-free decoding is superior because it is invariant to pθs’ phase relative to θ. Shaded regions indicate standard error of the mean of 10 simulated sessions.

We next modeled noise in the phase coupling between the theta carrier and place cells, either by perturbing neurons’ phase independently (“private noise”, Fig. 2 middle) or together (“shared noise”, Fig. 2 bottom). Private noise changes the phase relationships between neurons, and consequently, between the LFPs measured at different electrodes. This reduces the performance of both carrier-based and carrier-free methods, both of which rely on the fidelity of the multi-electrode patterns that define pθ. Shared noise, on the other hand, does not affect the phase offsets between place cells or electrodes, only their phase relative to the carrier. As carrier-based decoding is sensitive to this relative phase, it is compromised by shared noise. In contrast, because carrier-free decoding is insensitive to pθs’ phase relative to the carrier, it is robust to the presence of shared noise (Fig. 2F). In light of this finding, we devised a method for carrier-free decoding of physiological data and tested whether it improves our ability to recover information about position when theta is weak.

### A shallow neural network for carrier-free decoding

To learn pθs without using a carrier, we designed TIMBRE (Tracking Informative Multivariate Brain Rhythms Efficiently), a complex-valued artificial neural network (ANN) ^20^ with three layers (Fig. 3A; Extended Data Fig 3): 1) A complex-valued input layer that encodes the instantaneous phase and amplitude of theta at each electrode. This representation is equivalent to that used for carrier-based decoding; 2) A hidden layer, each node of which learns a unique complex-valued projection of the input that identifies a unique place-tuned pθ oscillation. The output of each node is a complex number; by calculating its magnitude and applying a ‘softmax’ function, the layer represents the probability that each pθ oscillation is present; 3) An output layer, each node of which predicts the probability that the rat occupies a particular behavioral state (e.g., maze arm) based on which pθ’s are present. TIMBRE’s design allows it to learn behaviorally-tuned oscillatory patterns from physiological data without the need of a carrier.

**Fig. 3.**
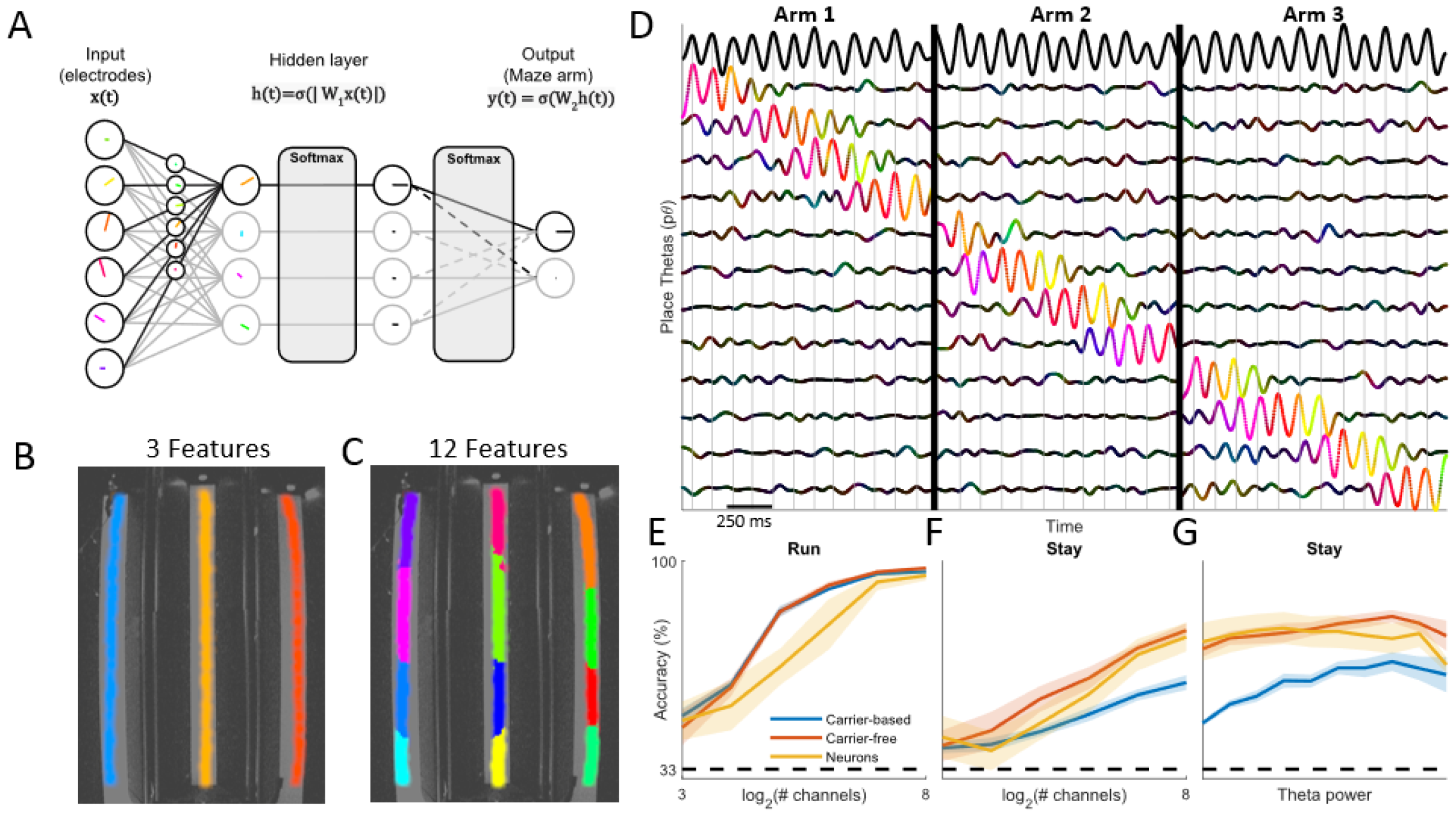
Carrier-free decoding of position from LFPs. **A)** TIMBRE architecture, consisting of a hidden layer that takes a complex-valued linear projection of the inputs. The absolute value of this projection is transformed using a softmax operation on the hidden layer. The hidden layer activations are projected onto an output layer that predicts which maze arm is occupied. **B)** When the hidden layer contains 3 nodes, each node learns to represent a single arm. **C)** When the hidden layer contains 12 nodes, each track is tiled uniformly by a subset of the nodes. **D)** During running, pθ’s (colored lines, network with 12 hidden nodes) activate sequentially and undergo phase precession relative to θ (top black line). pθ color indicates phase relative to θ and intensity indicates amplitude (as in Fig. **1C**). Vertical gray lines indicate θ peaks. Vertical black lines separate single runs on each of the 3 maze arms. **E)** During running, carrier-based, carrier-free, and spike decoders are similarly effective in predicting the rat’s arm occupancy. **F)** During staying, carrier-free and spike decoders outperform carrier-based decoding. **G)** Carrier-based decoding performance shows greater dependency on θ power than carrier-free or spike decoding. **E-G** depict mean ± s.e.m., n = 4 sessions from 3 rats.

TIMBRE’s hidden layer learns pθ oscillations whose place-tuned activations uniformly tile the environment - the larger the hidden layer, the finer the tiling (Fig. 3B-C, Extended Data Fig. 3). This is notable because the network is only trained to predict the maze arm being currently occupied, yet it learns a much more fine-grained representation of the rat’s position, a phenomenon known as feature learning ^21^. As the rat runs through the maze, pθs activate sequentially, exhibiting phase precession with respect to θ (Fig. 3D), as found in place cells ^2^. The hidden layers response is constrained to 1D manifolds, as would be expected from a neural code that represents the linear track (Extended Data Fig. 3). In contrast, stay periods do not reveal such a clear manifold structure, though early and late phases of the stay period show some separation (Extended Data Fig. 3).

During running, when θ is strong, carrier-based and carrier-free decoding comparably predict which maze arm is being occupied by the rat (Fig. 3E). In contrast, during stay periods, carrier-free decoding is significantly more accurate at predicting maze arm occupancy (Fig. 3F). Carrier-free decoding resembles spike rate-based decoding, in that their accuracy drops similarly from run to stay periods, (Fig. 3E-F), but remains consistent over the range of θ power observed during staying (Fig. 3G). For classifying position, TIMBRE outperforms two dimensionality reduction methods, CSP and ICA (Extended Data Fig. 3), both of which are commonly used to identify informative oscillations in multi-channel EEG and LFP recordings ^3,22,23^.

### pTheta and theta reflect distinct network components

We next compare pθ with θ, the high-variance oscillatory component identified using PCA and previously used as a carrier. When we reconstruct the multi-electrode LFPs using both θ and pθ, we find that θ explains >80% of the observed variance, while pθ accounts for <3% of the signal. In contrast, only pθ carries information about the animal’s position (Fig. 4A). During running, pθ oscillates at a higher frequency than θ; no such difference is seen during stillness (Fig. 4B) ^24^. The differences between θ and pθ parallel those observed between interneurons and pyramidal cells in the hippocampus: the former have higher firing rates that are relatively consistent across the maze, while the latter are strongly position-modulated, remaining silent across much of the environment (Fig. 4C). Furthermore, pyramidal cells oscillate at a higher frequency than interneurons during running (Fig. 4D) ^25^. On average, we find that interneuron activity is better predicted by θ, while pyramidal cell activity is better predicted by pθ both during running and staying (Fig. 4E), even after accounting for the differences in place tuning among these constituents.

**Fig. 4.**
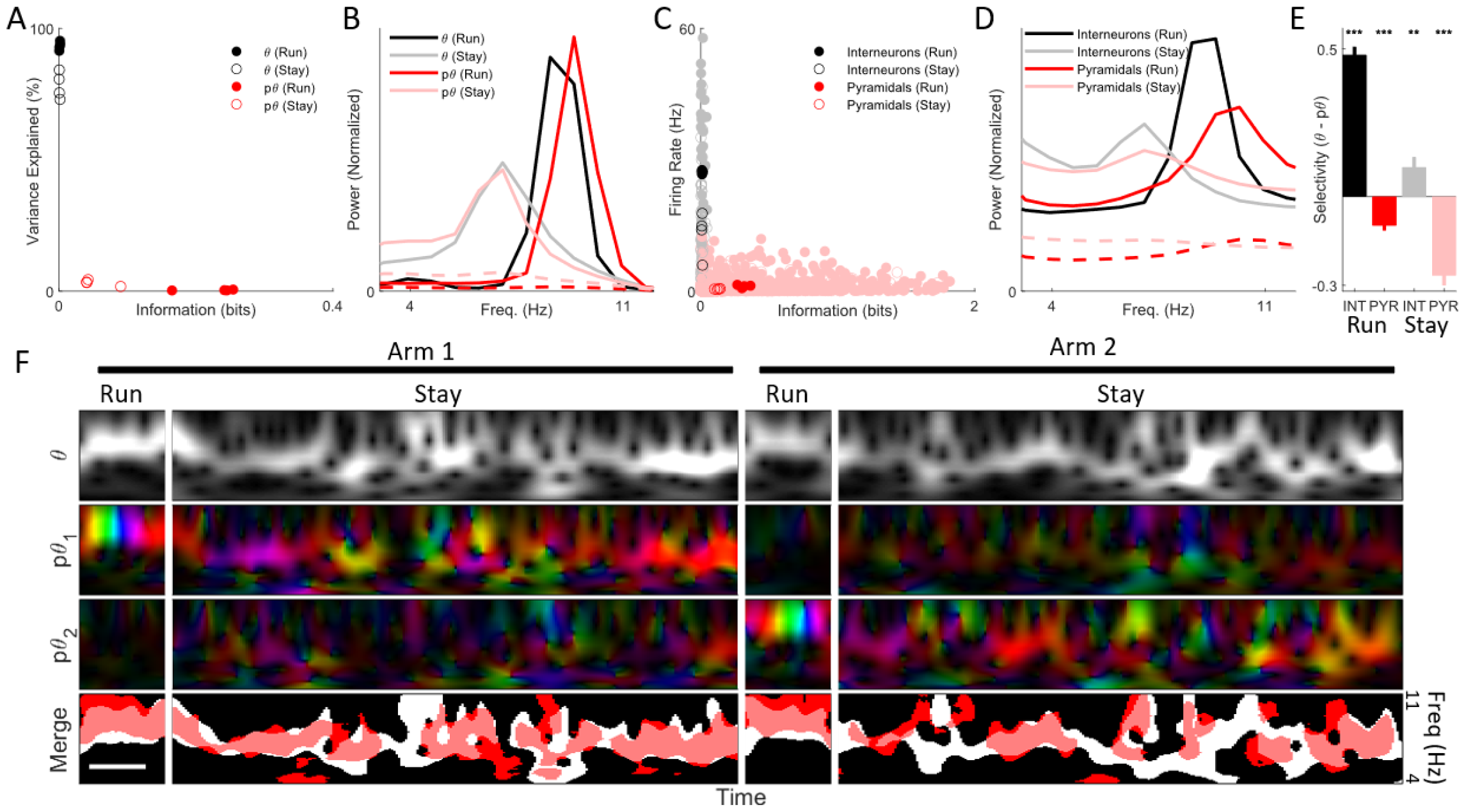
Physiological properties of two theta rhythms. **A)** θ accounts for most of the variance of the multichannel LFP (93% run, 78% stay), but carries no information about animal location. pθ explains little of the multichannel LFP’s variance (.41% run, 3% stay), but carries more information about animal location. Individual dots indicate variance and information during running (filled) and staying (empty) for 4 sessions. **B)** During running, pθ oscillates at a higher frequency than θ due to phase precession. During staying, the peak frequencies of pθ and θ overlap. **C)** Interneurons exhibit higher firing rates but are minimally modulated by position. Pyramidal cells exhibit lower firing rates but are strongly modulated by position. **D)** During running, pyramidal cells show higher peak frequency than interneurons. During staying, the peak frequencies of the two cell types overlap. **E)** Interneuron activity is better predicted by θ, while pyramidal cell activity is better predicted by pθ. ***p<.001, **p <.01 that the mean selectivity of each class of neurons is θ. Error bars depict mean ± s.e.m., n = 132/443 (INT/PYR run); 129/172 (INT/PYR stay) from 3 rats over 4 sessions. **F)** θ (top row) and pθ (2nd + 3rd rows) for two arms. pθi and pθ2 correspond to oscillations that are active in arms 1 and 2 respectively; colors indicate phase relative to θ. During running, the relative phase changes due to phase precession. During staying, the relative phase drifts despite unchanging position. (Bottom row) Regions where θ and pθ power exceed threshold are indicated in white and red respectively. During running, pθ shows a consistently higher frequency than θ during staying, both oscillations show drift in amplitude and frequency that partially overlap. For **B** and **D**, red solid (dashed) lines indicate pθ and place cell spectra for trials in which the rat was in the signals’ preferred (nonpreferred) arm.

Observing θ and pθ on single trials reveals their complex and somewhat distinct dynamics. As the rat runs across the track, the higher frequency of pθ results in phase precession relative to θ. During staying, θ and pθ show a gradual drift in their relative phase and partial overlap in their power across frequency and time (Fig. 4F).

### Carrier-free decoding in the open field

We evaluated TIMBRE’s ability to extract information about position from theta oscillations during foraging in the open field. Because foraging involves a larger variety of movements, and because theta rhythm generation is sensitive to movement, it offers a test of the robustness of LFP-based decoding. When trained using weak supervision (Extended Data Fig. 5), we find that the pθ components learned by TIMBRE’s hidden layer are localized both in x-y position and heading direction (Fig. 5A). In contrast, individual place cells, while localized in position, often lack orientation selectivity (Fig. 5B). Each learned pθ component shows phase precession relative to θ (Fig. 5A).

**Fig. 5.**
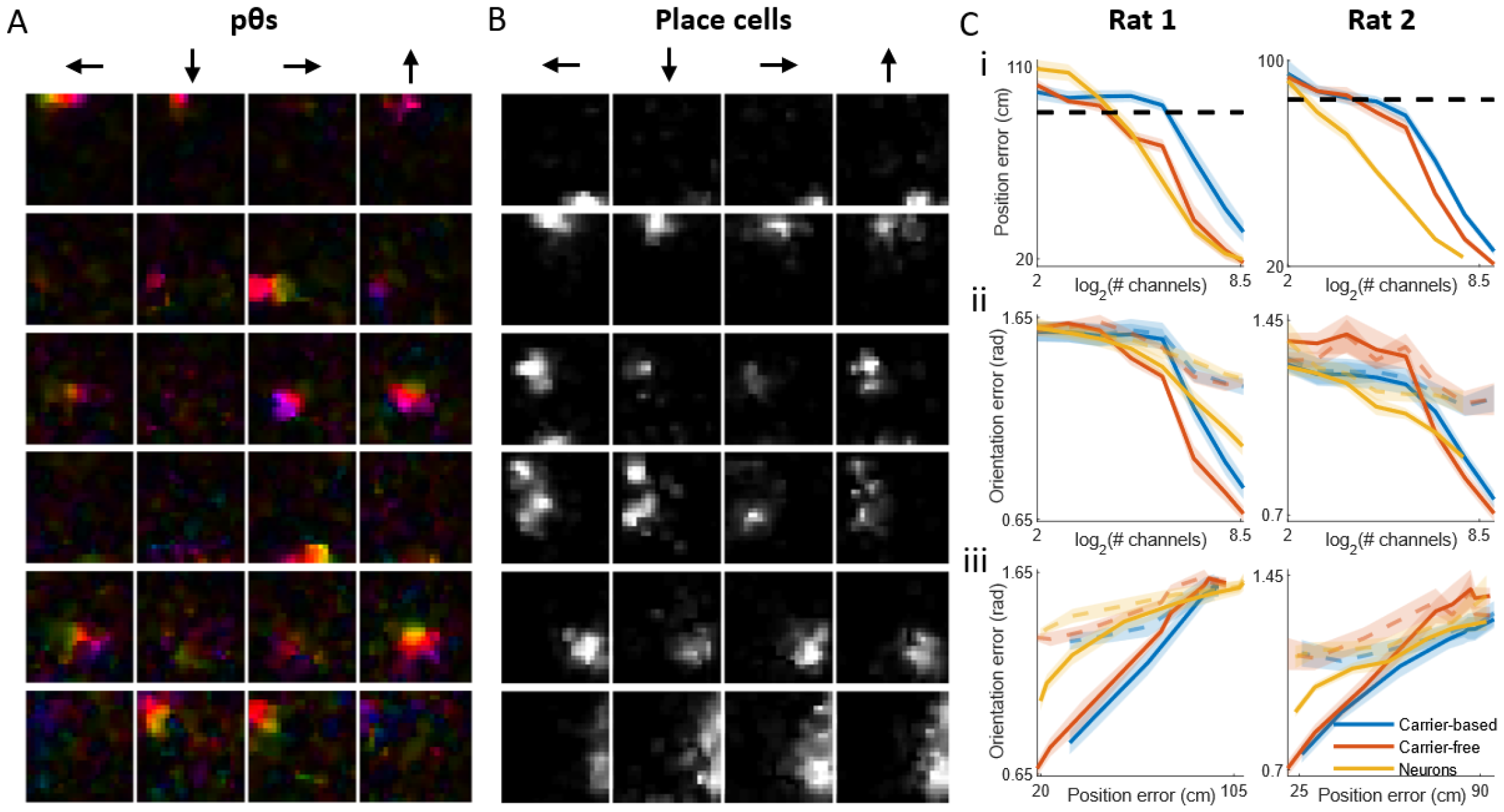
Decoding LFPs during open field foraging. **A)** Responses of 6 hidden nodes of TIMBRE as a function of rat position and orientation in the open field. Colors represent the phase of each pθ relative to θ. **B)** Responses of 6 pyramidal cells as a function of position and orientation in the open field. **C)** Mean error of carrier-based, carrier-free, and spike decoders in the open field for two rats, i) Position error as a function of # channels or neurons; ii) Orientation error as a function of # channels or neurons; iii) Orientation error as a function of position error. For i) black dashed line indicates chance-level performance; for ii and iii) colored lines indicate chance-level performance of orientation decoding achievable by position decoders, accounting for position-dependent orientation preference of each rat. Shaded regions show mean ± s.e.m. of test accuracy, n = 5 (4) folds for rat 1 (2). For Rat 2, one fold was excluded due to distributional shift (see methods).

We compared carrier-based, carrier-free, and spike-based decoding of position and orientation in the open field (Fig. 5C, Extended Data Fig 5). All decoding methods improved as the number of electrodes or temporal smoothing increased; with carrier-free decoding outperforming carrier-based decoding.

## Discussion

Interpreting LFPs is challenging due to their complex relationship to neural activity and poor spatio-temporal localization ^26^. One exception is the hippocampal theta rhythm, which appears to serve as a canonical brain clock coordinating sequential spiking in place cells ^27^. We previously showed that theta phase could provide a reference for reading out phase-amplitude patterns in LFPs ^3^. However, under behavioral conditions where theta weakens, it becomes unreliable as a reference. Here, we show that phase-amplitude patterns can directly be read out from LFPs without recourse to a theta clock. This finding suggests that place-encoding neural networks maintain a phase-position code, at least partially independent of the classically observed theta rhythm ^28^.

### pTheta reports place cell assemblies across behaviors

During running, place cell spikes exhibit phase precession relative to theta. We find analogous phase precession in pθ (Fig. 3D, 4B, 4F, 5A). Because pθ and θ are tightly coordinated during running, carrier-free and carrier-based models predict position equally well (Fig 1C-E left, 2E, 3B). Consequently, during running, we find no evidence that place cell assemblies oscillate independent of theta. Carrier-free decoding with TIMBRE reveals place-tuned rhythmic components that trace a 1D manifold (Extended Data Fig. 3), as expected from coherent place cell populations. We find different hidden layer sizes to reveal different degrees of place field localization (Extended Data Fig. 3), reminiscent of the multi-scale nature of the place code ^29^.

When locomotion stops, a weak and irregular ‘Type II’ theta is observed (Fig. 1F; Fig 4B, F) ^4^. In this condition, carrier-based decoding suffers (Fig. 1G). One possible explanation for this drop is that when theta is weak, it can no longer synchronize place cells and consequently fails to generate informative patterns in the LFP. Alternatively, place cells could generate coordinated activity independent of theta. Consistent with the second explanation, a carrier-free model better predicts position across a range of θ power, similar to spike-based decoding (Fig. 3F-G). Unlike during running, during stay periods decoding improves by using multiple hidden nodes to represent each location. This suggests that a drop in θ power leads to more variability in place cell population activity, and by extension, in position-tuned LFP patterns. While the time-averaged power spectra of pθ and θ overlap (Fig 4B), the two rhythms show distinct time-varying activity (Fig 4F). During both run and stay periods pθ and θ predict pyramidal cell and interneuron activity, respectively. Notably, the association between pθ and pyramidal cells extends beyond their place selectivity (methods), suggesting that pθ is reporting noise correlations among place cells.

The open field provides a third behavioral context for examining pθ. Unlike the linear track, in the open field place cells show localized place fields that are invariant to the rat’s direction of motion (Fig. 5B). While only trained to classify position, TIMBRE’s hidden layer reveals pθs that are selective for both position and orientation and that phase-precess in the direction of motion (Fig. 5A). Given that place cells exhibit phase precession in the open field ^5^, we would expect these cells to activate in an orientation-dependent order. Consequently, movement in each orientation would evoke distinct phase patterns in the LFP. The orientation selectivity of pθ thus provides further evidence that it reports sequential activity within place cell assemblies.

### TIMBRE, a neural network for identifying information-bearing brain rhythms

A further contribution of this work is TIMBRE, a simple and interpretable neural network that extracts oscillations that are predictive of behavior. TIMBRE achieves carrier-free decoding by learning oscillatory patterns whose amplitudes vary in a behavior-dependent manner. We found that using a softmax transfer function encourages competition among nodes in the hidden layer so that they learn features that tile the behavioral space. When using carrier-free and spike-based decoders as baselines, TIMBRE appears to be more successful in decoding position in the maze than in the open field. One potential explanation for this difference is that the maze involves a more stereotyped behavior in which each position is heavily sampled. In contrast, foraging in the open field invokes a more diverse behavioral repertoire, presumably evoking a wider array of LFP patterns. At the same time, the open field is less thoroughly sampled, limiting the training data for each hidden node. Finally, while our recordings had at most 255 channels, we see evidence that decoding will continue to improve with more electrodes, both in the maze (Fig. 3F) and the open field (Fig. 5C).

TIMBRE complements many existing computational approaches that have been developed to decode and interpret multi-channel brain wave data. Like complex-valued ICA (cICA) ^30^, it learns a complex-valued linear projection of the data that maximizes a particular objective. When applied to the theta rhythm during running, these algorithms produce similar results (compare Fig. 3C to Fig. 4A of ^3^). Because cICA is fully unsupervised, many of its learned features may reflect artifacts unrelated to brain function ^31^. In contrast, TIMBRE aims to discover a feature space that is maximally informative about a behavior of interest. Another related approach is Common Spatial Patterns (CSP), which is used to identify oscillatory subspaces whose variance changes maximally across conditions ^32^. We find CSP to be inadequate for decoding and interpreting theta LFPs (Extended Data Fig 3D), suggesting its orthogonality and variance constraints are unsuitable for revealing low-variance multi-electrode oscillatory patterns that occupy a manifold (Extended Data Fig. 3E). Another parallel is found in shallow convolutional networks that have been developed to detect behavior-encoding waveforms from EEG data ^33,34^. In particular, EEGNet reduces the number of parameters by enforcing spatiotemporal separability within its learned features. However, TIMBRE has the advantage of being able to detect phase offsets across electrodes, invariant of timescale. Because place cells preserve phase relationships despite changes in theta frequency ^35^, TIMBRE can detect pθ even as its center frequency shifts, as is seen during periods of stillness (Fig. 4F). We therefore believe that TIMBRE is ideal for multi-electrode recordings in brain regions where neurons are phase-coupled and respond to a measurable behavior or condition.

### Open questions

This work raises several questions for future research. First, what are the neural underpinnings of pθ and θ, respectively? One possibility is that these correspond to the two components in the ‘dual oscillator’ model ^2^, with θ reflecting pacemaker input from the medial septum and pθ potentially signaling the collective activity of place cells and/or inputs from grid cells within the entorhinal cortex ^36^ (Extended Data Fig. 6). While it remains possible that both rhythms arise from a single network ^24^, we believe that the distinction between pθ and θ is meaningful, in that they selectively correlate with pyramidal cells and interneurons respectively (Fig. 4E). Second, what are the functional implications of the greater variability observed in theta during stay periods? On the one hand, this variability could result from a low-fidelity representation of self-position due to reduced navigational demands during stay periods. On the other hand, the variability in phase coupling we observe (Fig. 4F) could result from a more varied behavioral repertoire during staying, potentially indicating shifting contributions of memory encoding and retrieval ^37^. In conjunction with mechanistic and behavioral interventions, TIMBRE could help address such questions by providing a mesoscopic view of neural codes.

---

**Extended Data Fig. 2.**
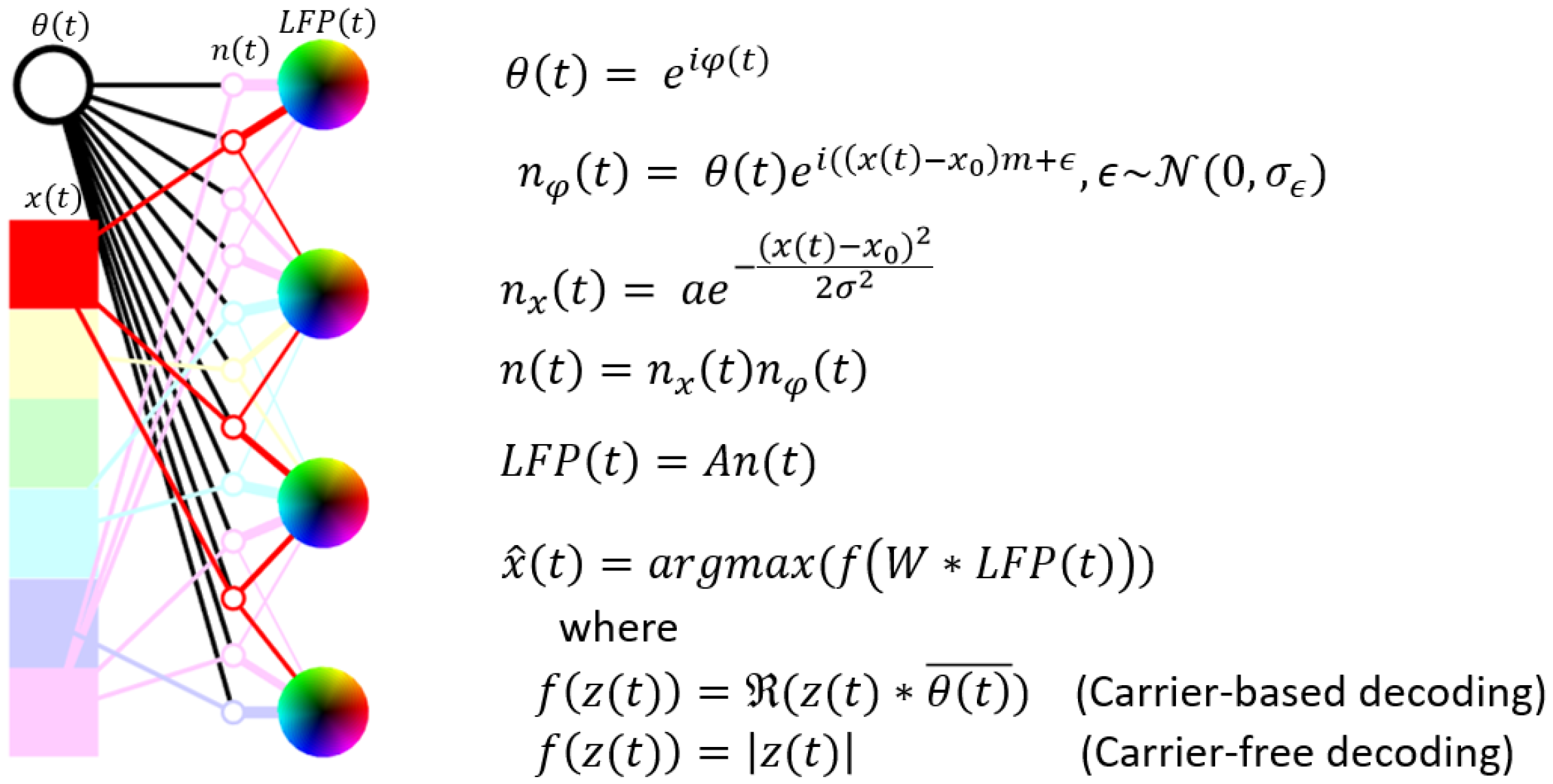
Model of LFP generation. Neurons receive position (x(t)) and theta (θ(t)) inputs, which modulate position-dependent (n_x_(t)) and theta-dependent (n_φ_(t)) components of activity respectively. Neuronal responses (n(t)) of place cells are modeled as a product of the position (n_x_(t)) and oscillatory (n_φ_(t)) components, each of which is complex-valued. Place cell responses are summated using a random mixing matrix A to produce measurements at the electrodes (LFP(t)). Multiplying LFP(t) by the appropriate demixing matrix W results in position-tuned oscillations. For carrier-based decoding, these oscillations are demodulated using θ(t), and position is estimated based on the oscillation with the largest real component. For carrier-free decoding, position is estimated based on the oscillation with the largest amplitude.

**Extended Data Fig. 3.**
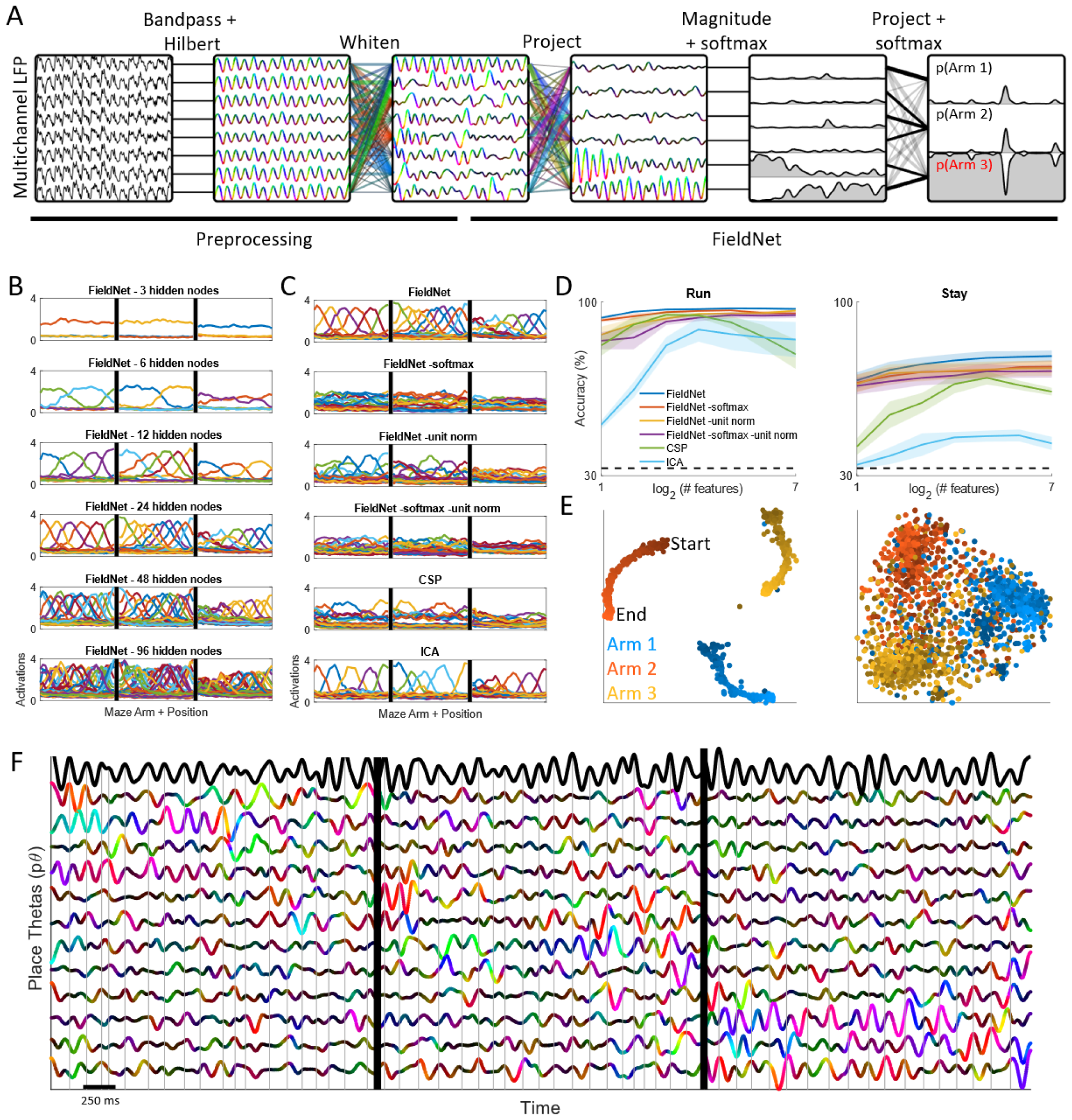
TIMBRE versus alternatives for identifying informative oscillatory subspaces. **A)** Pipeline for carrier-free decoding of LFPs. Broadband LFP is bandpass filtered (2-10 Hz) and downsampled and whitened using SVD before being fed into TIMBRE, which learns a collection of oscillations that are informative about behavioral state (in this case, maze arm occupancy). **B)** Average response magnitudes of hidden nodes as a function of maze position, for increasing hidden layer sizes. Different colors indicate activations of different hidden nodes. **C)** Feature activations for different versions of TIMBRE (with or without softmax and unit norm), as well as two alternative algorithms, CSP and cICA. In each case, 24 features are depicted. **D)** Comparison of arm classification performance for algorithms shown in C during running (left) and staying (right), mean ± s.e.m., n = 4 sessions from 3 rats.. **E)** During running (left), hidden layer activity traces out 1D manifolds corresponding to the linear geometry of each maze arm. Darker shades correspond to earlier phases of the trial. During staying (right), no clear manifold structure is apparent; nonetheless earlier (darker) and later (lighter) phases of the trial appear to segregate. **F)** As with running (Fig. **3D**) pθ’s activate selectively in one arm; however, there is no clear sequential ordering of pθ activations.

**Extended Data Fig. 5.**
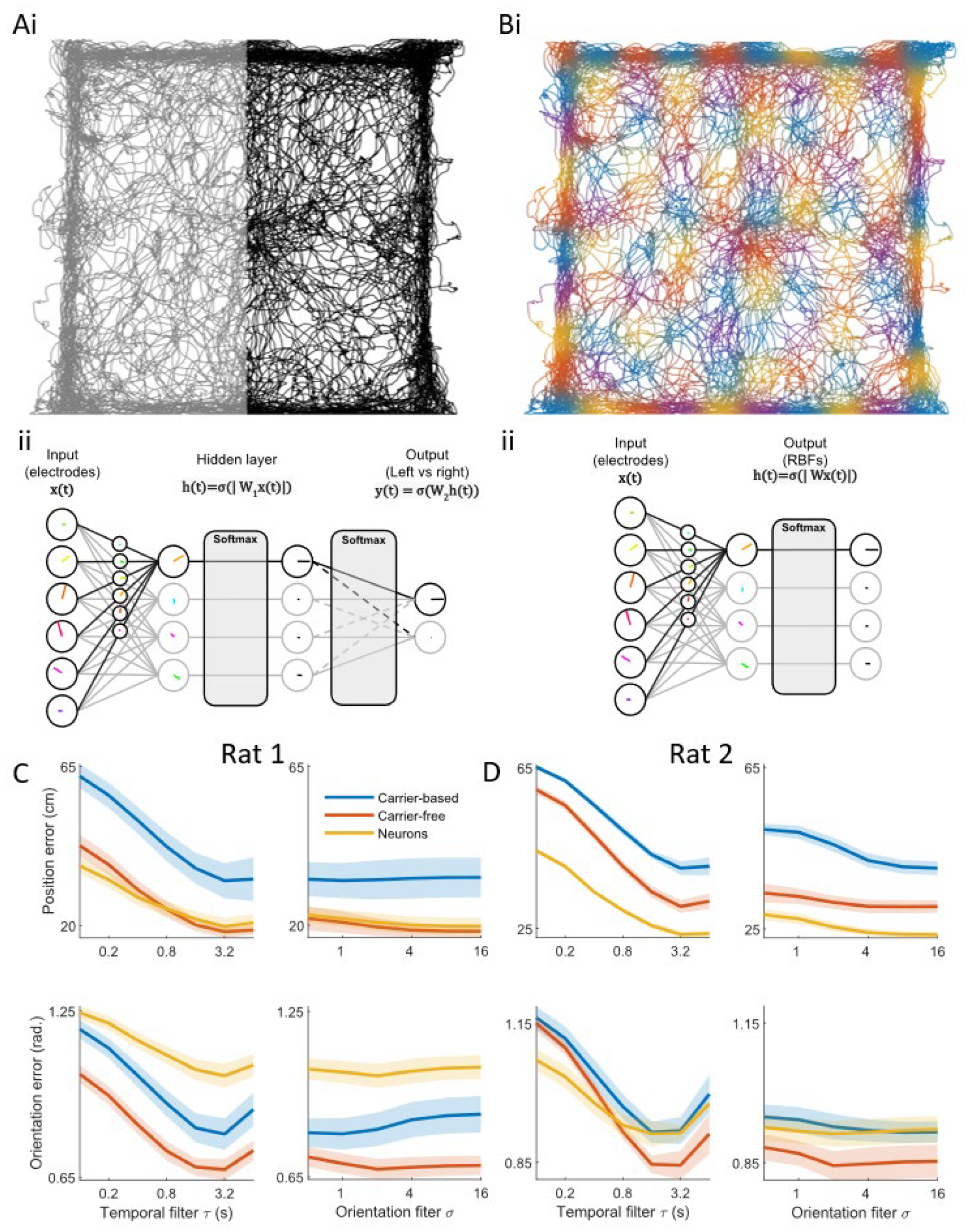
TIMBRE for representational learning or direct estimation. **A)** Representational learning, i) Positions are labeled in a binary manner by splitting the field into two. ii) the hidden layer learns a maximally informative collection of features to distinguish the two halves. **B)** Direct estimation. Positions are represented using a collection of radial basis functions (RBFs). RBF widths are depicted here more narrowly than what was used during training, for visual clarity, ii) TIMBRE learns a projection that predicts the activation of each RBF, without a hidden layer. **C)** and **D)** For rats 1 and 2 respectively, position (top) and orientation (bottom) error as a function of temporal filter τ (left) and orientation filter σ used to smoothen decoder output. Shaded regions show mean ± s.e.m. of test accuracy, n = 5 (4) folds for rat 1 (2). For rat 2, one fold was excluded due to distributional shift (see methods).

**Extended Data Fig. 6.**
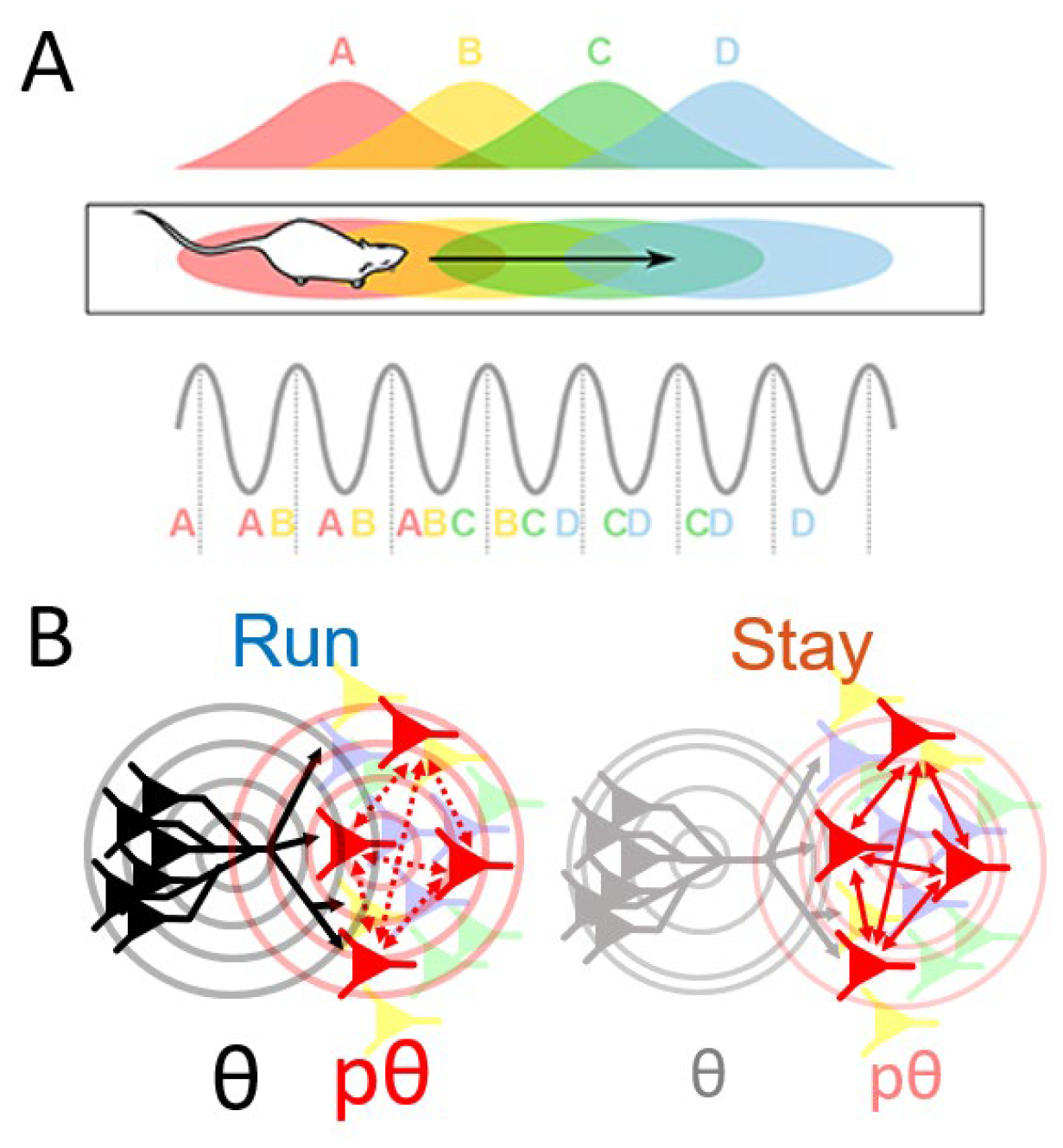
Proposed relationship between θ and pθ rhythms and hippocampal physiology. **A) A)** As a rat runs through its environment, the θ rhythm (gray wave) is consistently seen, while neurons fire action potentials (colored letters) at specific phases of the theta cycle in a sequence that is determined by the animal’s trajectory. Figure reproduced from 13. **B)** The θ rhythm, readily visible at individual recording sites, may arise from a population of neurons such as interneurons in the hippocampus and/or the medial septum. Conversely, the pθ rhythms, extracted by TIMBRE, are spatiotemporal waveforms whose shape is determined by the cunent location of the rat in its environment. We think that p0s arise from coherent activity in large populations of place-tuned cells. During periods of running (left), θ and pθ rhythms are robust and in sync. In this condition one cannot tell whether pθ arises due to a common θ input (black arrows), direct coupling among place-tuned neurons (red dashed arrows), or both. During stay periods (right), θ and pθ are less rhythmic and more out of sync with each other. This decoupling of the rhythms suggests that place-tuned cell assemblies can resonate autonomously without θ input (red solid arrows). While the cells generating θ and pθ are shown here as anatomically segregated, they may be interspersed.

## Methods

### Subjects

Adult male Long-Evans rats were used for experiments. All procedures were performed according to the Janelia Research Campus Institutional Animal Care and Use Committee guidelines on animal welfare.

### Three Arm Delayed Sequence Task

The Three Arm Delayed Sequence task (TADS) requires the rat to visit maze arms for reward in the following order: left, center, right, and center. The rat starts each trial in the delay area and has to run in a wheel in a fixed direction continuously for eight seconds over a set speed threshold to activate all three maze arm doors to open. The 2.13m-long maze (Fig. 1B) was designed by Eva Pastalkova and Tanya Tabachnik in the Instrument Design & Fabrication facility at Janelia Research Campus. The design and operation of the maze is described in ^38^.

### Recording neural activity

We performed chronic extracellular recording in freely behaving rats performing the spatial memory task. Chronic extracellular recording of neural activity was performed using silicon probes from Neuronexus (Buzsaki 64 and BuzsakiSP 64 probes). 3d printed ring, external faraday cage, and nanodrives were from Ronal Tools (http://www.ronal.com/). Ring was designed by Eva Pastalkova and Yingxue Wang. Implant procedure was by Eva Pastalkova. Faraday cage was designed by Brian and Andy Lustig.

Neural signals and tracking of position was carried out using the Amplipex recording system (http://www.amplipex.com/). Importantly the Amplipex system is integrated with Intan multiplexing headstages (http://www.intantech.com/index.html) which permit recording from large numbers of channels over a thin light cable to minimize interference with behavior in the freely moving rats. Commutators for recording were either custom from Dragonfly (www.dragonflyinc.com) or Doric Lenses (ERJ 12 HDMI-B2). Neural activity was recorded at 20 kHz.

Neural recordings were stored as .dat files. Behavioral location within the maze was recorded by tracking of LEDs on the rat’s implant at 30Hz using a webcam integrated with the Amplipex system. A sync pulse from the maze behavioral control system was recorded on the Amplipex system in order to guarantee accurate syncing of behavioral information and neural activity recorded on Amplipex. For LFPs, recordings were downsampled to 1250 Hz and saved as .lfp files.

### Spike sorting and interneuron classification

The NDManager and Klusters suite was used for analysis as well as MATLAB. Spike detection was performed using high pass filtered signals and threshold crossings above the mean + 1.5 SD. Then spike waveforms were stored and sorted using the KlustaKwik suite. After automatic sorting, clustering was further refined manually using Klusters ^39^. Each isolated unit was assigned an interneuron score by BL on a 6-point scale based on 3 criteria: 1) the asymmetry of the spike waveform: for interneurons, the hyperpolarization peak is larger in amplitude than the depolarization peak; 2) the spike width: interneurons have narrower spikes; 3) the firing rate: interneurons have higher firing rates. Neurons scoring a ‘0’ or ‘5’ (i.e., the extremes of the 6-point scale) were defined as pyramidal cells and interneurons respectively for subsequent analysis (Fig. 4).

### LFP preprocessing

LFP recordings were 1) downsampled from 1250 Hz to 25 Hz (using Matlab’s ‘decimate’ function, which applies a 10 Hz, 8th-order Chebyshev Type I low-pass filter prior to downsampling to prevent aliasing). 2) high-pass filtered (2 Hz, 2nd-order Butterworth filter, zero-phase filtering performed using Matlab’s ‘filtfilt’ function). 3) Hilbert-transformed to produce a complex-valued signal (using Matlab’s ‘hilbert’ function) that represented the instantaneous amplitude and phase. Additional pre-processing for carrier-based and carrier-free decoding is described in the next 2 paragraphs respectively.

For carrier-based decoding, the first PC (the “carrier” or θ) of the complex-valued LFP was identified using matlab’s ‘svd’ function. The phase of the carrier was subtracted from the phase of each channel to produce a demodulated LFP

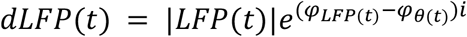

Finally, the signal was converted from a *T* x *N* complex-valued matrix to a *T* x *2N* real-valued matrix (i.e., with real and imaginary components concatenated).

For carrier-free decoding, the LFP was whitened using matlab’s ‘svd’ function, and rescaled by dividing each dimension of the data *X*(denoted *X*_*I*_) by

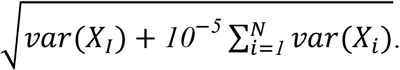

Analogous to ridge regression, this transformation rescales the data but reduces the contribution of the smallest whitened dimensions, which tend to be more noisy.

### LFP analysis in the maze

#### Identifying ‘run’ and ‘stay’ periods

‘Run’ periods were defined as times when the rat was in one of the three maze arms approaching a reward port. Other periods, including when the rat was running away from the reward port, were discarded as these periods had less consistent running behavior (i.e., with frequent stops) across trials and sessions. ‘Stay’ periods were defined as times when the rat was positioned near a reward port for the purpose of consuming water rewards. Unrewarded trials were discarded as these consisted of very brief stay periods. ‘Run’ and ‘Stay’ periods were analyzed separately for all subsequent analyses.

#### Data selection for cross-validation

Data from each trial was randomly assigned to one of 5 folds, ensuring that data used for model training and testing were separated in time. Since rats traversed the center arm twice as often, and stayed at some reward ports longer than others, data was sub-sampled so that each arm was equally represented for the purpose of model training (i.e. so that the model would not be biased to select the most occupied arm), and such that all 5 folds had equal amounts of training data. 100 random fold assignments were tested, and the assignment that produced the largest amount of training data, given the above subsampling procedure, was chosen.

#### Probabilistic decoding of arm occupancy (Fig. 1D)

L1-regularized logistic regression ([liblinear](https://www.csie.ntu.edu.tw/%7Ecjlin/liblinear/)^40^) was used to predict from the demodulated LFP which arm was being occupied by the rat. The cost parameter was set to its default value of 1. The probability of each arm being occupied was determined on a held-out trial using liblinear’s ‘predict’ function.

#### Carrier-based decoding (Fig. 1F and 3E-G)

An L1-regularized logistic classifier was trained using either ‘run’ or ‘stay’ data to predict from the demodulated LFP which maze arm the rat occupied. For running, each track was divided into four sections, for a total of 12 classes. This accounted for the changing place code along the length of the track. The classifier was evaluated on held-out data. Accuracy was defined as the percentage of samples whose predicted category (12 for run, 3 for stay) was found in the arm currently occupied by the rat. To evaluate LFP (spike) decoders’ dependency on the number of electrodes (neurons), the first n channels were used as inputs to the decoder, where n sampled powers of 2 from 8 to 256.

#### Carrier-free decoding (TIMBRE)

A complex-valued neural network was implemented in Keras using the [complexnn](https://github.com/JesperDramsch/keras-complex) package ^20^. The network has one hidden layer, with both the real and imaginary components of the weights of each node constrained such that their L2 norm equals 1. The output of this layer was transformed using the absolute value function, thus discarding the phase, followed by a ‘softmax’ operation. This operation was found to induce competition between hidden nodes, encouraging them to learn different features. The hidden layer was fed into an output layer with 3 nodes (one for each maze arm) with a softmax operation that estimates the probability of each arm’s occupancy. The network was trained using Adam optimizer, learning rate = .001, and categorical cross-entropy loss function. Training a network with the combination of cross-entropy loss and softmax has been shown to maximize mutual information between the inputs and the outputs ^41^. Training was stopped at 100 epochs, or when the test loss increased (i.e. ‘early stopping’), whichever came first (usually the latter condition, in practice).

#### Spike-based decoding

A TxN matrix was defined, each entry containing the number of spikes fired by each neuron within a 25-Hz time window. The columns of this matrix were low-pass filtered (3 Hz, 8th-order Butterworth filter, zero-phase filtering performed using Matlab’s ‘filtfilt’ function). The frequency was chosen so that spike decoding was roughly the same accuracy as carrier-free decoding during running (lower cutoff frequencies resulted in smoother signals that provided better decoding). Each neuron’s response was normalized by its standard deviation calculated within ‘run’ or ‘stay’ periods. Logistic regression was then performed analogously to carrier-based decoding, described above.

#### pθ rendering

The preprocessed LFP was multiplied by the conjugate of the hidden layer’s weights, producing one pθ oscillation per hidden node. *Spatial rendering:* the magnitudes of pθ activations were binned by 2-d position within the track and averaged per bin using matlab’s accumarray function. Locations were binned into square bins, such that there were 200 bins along the length of the track. Before rendering, spatial maps were smoothed with a gaussian kernel with σ = 2 bins. Each pixel was assigned a unique color corresponding to the most active pθ at that location, with the alpha channel set to the average activation for the pθ. Activations exceeding 1 were set to 1. *Temporal rendering:* pθs were sorted by their peak activation within the 3 arms. pθ phases were rotated so that they had θ average phase lag with respect to θ (as estimated using Matlab’s correlation function). Each pθ response was normalized, dividing by the 95th percentile of its response magnitude. Data from 3 trials, one per arm, were concatenated. Plots show real projection of data, with color hue indicating phase of the demodulated pθ (i.e., pθ phase relative to θ) and intensity indicating response amplitude (with black representing low-amplitude). *Manifold visualization:* The magnitudes of pθs across 1 session (from TIMBRE with 48 hidden nodes), for held-out data from either ‘Run’ or ‘Stay’ periods, were fed into matlab’s built-in tSNE function, using the ‘correlation’ distance metric. Each arm’s data was color coded using a separate color, with progressively brighter colors assigned to later parts of each trial. Data was 2x downsampled for the longer ‘Stay’ periods using Matlab’s ‘decimate’ function, to reduce run time.

#### Comparison to CSP and ICA

Víctor Martínez-Cagigal’s implementation of CSP ^42^ was downloaded from Matlab’s File Exchange. It was modified to handle multiple classes of data by, for each class, solving the generalized eigenvalue problem *C*_*i*_*x = λCx*, where *C*_*i*_ is the covariance matrix for complex-valued LFPs recorded when the rat was in arm *i*, whereas *C* was the covariance matrix for data collected in all arms. The top ‘n’ eigenvectors, as determined by their eigenvalues, were taken for each arm. For complex-valued ICA (Novey & Adali 2006), a vectorized version of the ACMNsym matlab function was used and the top ‘n’ components, as determined by the L2 norms of the components of the mixing matrix, were taken for each arm. For both CSP and ICA, the amplitudes of their activations were used to predict the current arm using L1-regularized logistic regression (as done with carrier-free decoding above) (Extended Data Fig. 3D), and averaged by position to generate spatial activation profiles (Extended Data Fig. 3C).

#### Power spectra

The Chronux package (chronux.org^43^) was used to generate power spectra. Specifically, since trials were of unequal length, a short time fourier transform (STFT) was taken on consecutive windows of data within the appropriate trial period (i.e., during run or stay periods) with the taper parameter set to [1/T, T, 1], where T = window size in seconds. The resulting power spectra were averaged across the session. *Fig. 1:* SVD was used to produce principal components of the LFP. The spectrum for the first PC was plotted during run and stay, and the spectra of the remaining PCs were added together to generate the power spectrum of the residuals. A T = 1.5 s window was used. Fig. 4: For θ, the spectrum of the first PC was taken. For pθ and place cells, the spectrum of each signal was taken separately in each arm. The ‘preferred arm’ for each signal was defined as the arm in which the signal had the largest standard deviation. The other two arms constituted the ‘non-preferred arms’. All spectra (across 3 pθ’s, or all place cells) were averaged together to generate the preferred arm (solid line) or non-preferred arm (dashed line) spectra. For each session, spectra (representing all arms for θ and interneurons, or preferred arms for pθ and place cells) were normalized so that their area summed to 1, prior to averaging across sessions. Spectra for non-preferred arms (for pθ and place cells) were scaled to maintain their power relative to the preferred arm spectra. Error bars were excluded for clarity.

#### Wavelet transform (Fig. 4F)

A period of one trial from each of two arms was selected (1.52 sec for ‘Run’ and 10 sec for ‘stay’). A continuous wavelet transform was performed on the real component of θ or pθ (derived from TIMBRE trained with 1 hidden node per maze arm) using matlab function cwt, with default choices for wavelet (the complex-valued Morse wavelet) and bandwidth parameter (60) with 8 voices per octave and frequency range 2 Hz to 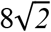 Hz. The resulting complex-valued signals were demodulated using the phase of θ, resulting in a zero-phase signal for θ and a slowly-varying phase for pθ (either due to phase precession during running or phase drift during staying). The matlab function scatteredInterpolant was used to transform the logarithmic frequency axis used by cwt to a linear one. Each signal was normalized by its standard deviation during run or stay periods, across all arms (for θ) or within the active arm (for pθ). For rendering, values of .9 or greater were set to maximum intensity, and a threshold of .4 was used to define active regions for the ‘merge’ image (bottom panel).

#### LFP variance explained by θ vs pθ (Fig. 4A)

For either the still or the run periods within each session, linear regression was performed to predict the multi-electrode LFP (bandpass-filtered and Hilbert transformed as described in **LFP Preprocessing**) from θ and 3 pθs (learned by TIMBRE with 3 hidden nodes). The samples used for training this model were the same as those used to estimate θ and pθ. Using samples from the test set, the variance of the reconstruction was calculated for each of the four regressors (θ and 3 pθs). This variance was normalized by the total variance of the LFP, and the variance explained by the 3 pθs was summed to produce a single number.

There were two important choices that influence our estimate of the fraction of the multi-channel LFP explained by pθ. First, we predicted the LFP jointly from θ and pθ, allowing θ to ‘explain away’ variance that would otherwise be captured by pθ. We did this because the LFP is so highly correlated, that any random projection of the data would appear to explain a substantial portion of the LFP. Second, we trained TIMBRE with 3, rather than more, nodes. Since pθs are projections of the LFP, larger numbers of pθs tend to overfit the LFP, fitting “noise” in addition to the position-encoding components of the LFP. Fundamentally, the finding that pθ and θ respectively reflect high-information and high-variance components of the LFP parallels our earlier finding that variance is concentrated in few dimensions but information is distributed across the full dimensionality of the LFP (Fig. 2C of ^3^).

#### Neuronal activity explained by θ vs pθ (Fig. 4E)

For each neuron, spikes were binned at 25 Hz sampling rate. The resulting vector was 1) low-pass filtered (10 Hz, 8th-order Butterworth filter, zero-phase filtering performed using Matlab’s ‘filtfilt’ function); 2) high-pass filtered (2 Hz, 2nd-order Butterworth filter using ‘filtfilt’); 3) Hilbert-transformed to produce a complex-valued signal (using Matlab’s ‘hilbert’ function). This produced a signal analogous to the preprocessed LFP (see above). For each session, TIMBRE was trained using 3 hidden nodes, resulting in one pθ that activated in each maze arm. Data was separated by arm and behavioral condition, resulting in six conditions per neuron. For each condition, the response of each active neuron (defined as those with firing rate greater than 2 Hz within the condition) were predicted using a linear model with θ and the active pθ as regressors. Each neuron’s selectivity *s* was calculated using the learned weights *w*_*θ*_ and *w*_*pθ*_ as 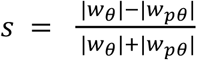. Because the regression was done separately for data from each arm, and because only one pθ was active in each arm, the selectivity of pyramidal cells for pθ was not due to their both being tuned to position (because pθ did not show position tuning within the condition). A two-tailed t-test was performed on the selectivity of each cell group (interneurons and pyramidal cells) within each behavioral condition (run vs stay) with the null hypothesis that the selectivity was not significantly different from 0.

#### Position information in LFPs and Neurons (Fig. 4A and C)

We calculated the information carried by LFPs or spike trains about position using the formula 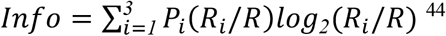. where *P*_*i*_ is the probability of occupying maze arm *i, R*_*i*_ is the average response (firing rate for neurons, amplitude for θ and pθ) in maze arm *i*, and *R* is the average response. For very sparse signals, this formula reports artificially high values ^44^; therefore, the information was calculated after randomly shifting the time series of maze arm occupancy relative to the time series of LFP/neuron responses, and averaged over 100 time-shifts. This randomized estimate was subtracted from the originally estimated information.

### LFP analysis in the Open Field

#### Outlier removal

Data from the session was divided into 5 consecutive, equally sized folds for the purposes of cross-validation. It was observed that there were occasional spikes in LFP power, potentially due to movement artifacts. These periods were removed by 1) running SVD on the complex-valued LFP data to determine the scores along the principal components; 2) z-scoring the scores; 3) summing the amplitudes of the scores; 4) z-scoring the sum and removing samples that exceeded a value of 2. Data was whitened after removal of these outliers from the training data. Excluding this removal step led to slight decreases in TIMBRE’s hidden-layer tuning and performance.

#### Visualizing position-orientation tuning (Fig. 4A and B)

TIMBRE was trained with a hidden layer of 256 nodes to classify the rat’s position within the left or right half of the open field (Extended Data Fig. 5). Compared to the linear track, the pθs were less strongly position-tuned. We therefore visualized the pθs in the open field after applying the softmax function to their amplitudes. This had the effect of suppressing the activity of hidden nodes in the locations where they were weakly activated. Finally, we demodulated the pθ’s by subtracting θ’s phase from the phase of each pθ. We then discretized the rats’ position into 24 bins along the x- and y-axis and into 4 orientations. We averaged the pθ or neuronal responses within each bin and applied a Gaussian spatial filter of radius = 1 bin for the responses within each of the 4 orientations. For visualizing neural place fields, we discarded the phases (leaving just the amplitudes) because spiking had a less consistent phase relative to θ, leading to noisier place fields when phase was kept. Example pθs (Fig. 4A) and neuronal (Fig. 4B) place-orientation fields were manually selected to highlight fields that were in the middle, rather than the edge, of the field. This is because the rat spent significant time running around the edges, and neuronal responses on the edge resembled those on the linear track, giving less insight into the nature of open-field responses.

#### Position and orientation decoding (Fig. 4C)

Position was rescaled so that it covered the range [0,1]. Position and orientation were represented by a 10 x 10 x 10 evenly spaced grid of radial basis functions using the formula 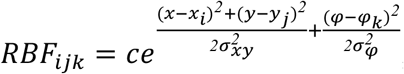. where *x,y*, and *φ* represent the animal’s 2d position and orientation; *x*_*i*_, *y*_*i*_, *and φ*_*k*_ represent the RBF’s center in position and orientation; σ_*xy*_ *= 1*/*9* and σ_*φ*_ *= 1*/*10* (values that are chosen so that RBF widths are concordant with their spacing; σ_*xy*_ and σ_*φ*_ are slightly different due to position being linear while orientation being circular), and *c* being a scaling constant such that the RBF responses sum to 1. For carrier-based decoding, the complex-valued LFP was demodulated, and its real and imaginary components were fed into a linear model with a softmax output, trained to predict RBF responses. For carrier-free decoding, the complex-valued LFP was projected into a layer whose amplitude was transformed using a softmax output, trained to predict RBF responses. KL divergence was used as the loss function for training. After training, RBF responses were smoothed in orientation and time by a range of values (indicated by σ and *τ* of Extended Data Fig. 5C/D), and the animal’s position and orientation were estimated to be the center of the maximally activated RBF. The average error between the estimated and true position and orientation were calculated. Using the training data, the orientation smoothing that led to the smallest training error (when *τ* = 6.4 seconds) was used to estimate position and orientation for test data. This was done separately for each rat and decoder (carrier-based, carrier-free, and neuron); in practice the choice of orientation smoothing had a small effect (Extended Data Fig. 5 C/D right panels). Chance-level position decoding was calculated by calculating the mean distance between the rat’s actual position and noise that was uniformly distributed over the range [0,1]. To evaluate LFP (spike) decoders’ dependency on the number of electrodes (neurons), the first n channels were used as inputs to the decoder, where n sampled powers of 2 from 8 to 256 (rat 1 had 192 electrodes, so this was the number of channels sampled instead of 256 for rat 1). Because a small number of neurons were orientation-selective, we derived more confident estimates on the decoding performance by averaging together the first ‘n’ and the last ‘n’ neurons. For rat 2, we noticed particularly poor test performance for the last fold, especially for carrier-free decoding. We traced this to a distributional shift in the eigenspectrum of the LFP. We therefore discarded this fold. Alternatively, carrier-free decoding could be rescued for this fold by normalizing the model’s outputs so that their average response to test data matched their average response to training data.

#### Position-conditioned orientation decoding (Fig. 4C, colored dashed lines)

Because the rat’s preferred orientation depended on its location in the open field, our ability to decode orientation could in principle be an artifact of decoding position. To account for this, we derived a model that estimated the rat’s orientation based only on its estimated position. First, we calculated the joint distribution of position and orientation for each rat by deriving RBFs for position only (10 x 10 x 1 grid) or orientation only (1 x 1 x 10 grid). We took the inner product of these two sets of RBFs to estimate the joint distribution of positions and orientations. Then, we estimated the animal’s orientation as the most likely orientation based on the rat’s estimated position.

### Decoding using model LFPs

#### Generation of multi-electrode LFPs

##### A population of place cells received two inputs

1) the position of an agent traversing a unit-length circular track in one direction, and 2) the theta rhythm, encoded as a complex-valued phasor with an amplitude of 1. The place fields of these cells were spaced uniformly along the track, with Gaussian tuning (*n*_*x*_ in Extended Data Fig. 2, with *a = 1* and σ *= e*^−*2*^). The theta-coupling of the neuron’s response *n*_*φ*_ exhibited phase precession at a slope specified by 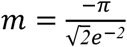 (see Extended Data Fig. 2). The position- and theta-dependent components *n*_*x*_ and *n*_*φ*_ were multiplied to generate a complex-valued response for each neuron. These responses were randomly mixed to generate a multi-electrode LFP by multiplying the neural responses with a matrix *A* of size *# electrodes* × *# neurons*. The real and imaginary components of *A* were sampled from a uniform distribution over the range [*0,1*] (sampling a 0-mean complex-valued Gaussian distribution led to weaker signals due to more phase cancellation between neurons). Noise was added to the phase of neurons by sampling from a normal distribution, either independently for each neuron (private noise), or identically for all neurons (shared noise). *Fig 2A-E:* 64 neurons were used (for visualization purposes, using few neurons produced visible phase differences across electrodes), with the agent running a single lap in 1.5 seconds, at a sampling rate of 800 Hz and theta rhythm of 8 Hz. A filter was applied (using Matlab’s gausswin function with window size = 16 electrodes and *α = 4*) to the columns of mixing matrix *A* so that the signal would change smoothly across electrodes, and a lowpass filter (Butterworth filter of order 4 and cutoff frequency *f*_*L*_) was applied to the noise so that it would change smoothly in time. For the shared noise condition, σ_*ϵ*_ *= 1* and *f*_*L*_ *= 8* Hz. For the private noise condition, σ_*ϵ*_ *= 2* and *f*_*L*_ *= 4* Hz. *Fig 2F (decoding):* 1000 neurons were used, sampling position and theta phase uniformly over the range [0,1] and [0,2*π*], respectively. Noise was sampled from a normal distribution with σ_*ϵ*_ *= 1*. Position was discretized into 25 equally-spaced bins for the purposes of decoding.

#### Decoding of simulated LFPs

The response of the neural population was calculated at each discretized location along the track, turning off the noise as well as the theta input. This response was then multiplied by the mixing matrix *A* to generate LFP matrix *X*. Demixing matrix *W* was calculated using the formula 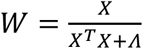, where *Λ = 10*^−*1*^ × *diag(diag(X*^*T*^*X*)). The demixing matrix *W* was multiplied by the demodulated LFP to generate a matrix of 25 pθ time series (one for each location along the track). For each sample, the entry with the pθ with the largest real (absolute) value indicated the estimated location for carrier-based (carrier-free) decoding (Extended Data Fig. 2).

## Data availability

Data is available at https://figshare.com/articles/dataset/Behavior_and_spiking_data_for_rats_running_a_3-arm_maze/24757638.

## Code availability

Code will be available at https://github.com/beatLaboratory/TIMBRE.

## Acknowledgements

We thank members of the Redwood Center and BeatLab for discussion. Funding was provided by NIH grant #1R01EB026955 and the Howard Hughes Medical Institute.

## Author contributions

The project was originally conceptualized by G.A. and F.S. The behavioral paradigm and data collection were developed by B.L., E.P. and A.L. Data was collected by B.L. TIMBRE, data analysis, and data visualization were developed by G.A. TIMBRE optimization and schematic was developed by S.A. Software repository was created by G.A. and S.A. The manuscript was written by G.A. and F.S. and edited and reviewed by all authors.

## Additional Information

**Correspondence and requests for materials** should be addressed to Gautam Agarwal or Friedrich Sommer.

## References

1. Herreras, O. Local Field Potentials: Myths and Misunderstandings. Front. Neural Circuits 10, (2016).

2. O’Keefe, J. & Recce, M. L. Phase relationship between hippocampal place units and the EEG theta rhythm. Hippocampus 3, 317–330 (1993).

3. Agarwal, G. et al. Spatially Distributed Local Fields in the Hippocampus Encode Rat Position. Science 344, 626–630 (2014).

4. Vanderwolf, C. H. Hippocampal electrical activity and voluntary movement in the rat. Electroencephalogr. Clin. Neurophysiol. 26, 407–418 (1969).

5. Jeewajee, A. et al. Theta phase precession of grid and place cell firing in open environments. Philos. Trans. R. Soc. Lond. B. Biol. Sci. 369, 20120532 (2014).

6. Buzsaki, G. Rhythms of the Brain. (Oxford University Press, 2006).

7. Bullock, T. H. Signals and signs in the nervous system: The dynamic anatomy of electrical activity is probably information-rich. Proc. Natl. Acad. Sci. 94, 1–6 (1997).

8. Gao, R. D. The (Epi)Phenomenal Oscillation, Spike, and LFP | mind from matter. https://www.rdgao.com/epiphenomenal-oscillations/.

9. Gray, C. M., König, P., Engel, A. K. & Singer, W. Oscillatory responses in cat visual cortex exhibit intercolumnar synchronization which reflects global stimulus properties. Nature 338, 334–337 (1989).

10. Frady, E. P. & Sommer, F. T. Robust computation with rhythmic spike patterns. Proc. Natl. Acad. Sci. 116, 18050–18059 (2019).

11. Akam, T. & Kullmann, D. M. Oscillatory multiplexing of population codes for selective communication in the mammalian brain. Nat. Rev. Neurosci. 15, 111–122 (2014).

12. Jonas, E. & Kording, K. P. Could a Neuroscientist Understand a Microprocessor? PLoS Comput. Biol. 13, e1005268 (2017).

13. Drieu, C. & Zugaro, M. Hippocampal Sequences During Exploration: Mechanisms and Functions. Front. Cell. Neurosci. 13, 232 (2019).

14. Eliav, T. et al. Nonoscillatory Phase Coding and Synchronization in the Bat Hippocampal Formation. Cell 175, 1119-1130.e15 (2018).

15. Dunn, S. L., Town, S. M., Bizley, J. K. & Bendor, D. Behaviourally modulated hippocampal theta oscillations in the ferret persist during both locomotion and immobility. Nat. Commun. 13, 5905 (2022).

16. Goyal, A. et al. Functionally distinct high and low theta oscillations in the human hippocampus. Nat. Commun. 11, 2469 (2020).

17. Oppenheim, A. V., Schafer, R. W. & Buck, J. R. Discrete-Time Signal Processing. (Prentice Hall, 1999).

18. Chadwick, A., van Rossum, M. C. & Nolan, M. F. Independent theta phase coding accounts for CA1 population sequences and enables flexible remapping. eLife 4, e03542 (2015).

19. McClain, K., Tingley, D., Heeger, D. J. & Buzsáki, G. Position–theta-phase model of hippocampal place cell activity applied to quantification of running speed modulation of firing rate. Proc. Natl. Acad. Sci. 116, 27035–27042 (2019).

20. Trabelsi, C. et al. Deep Complex Networks. in (2018).

21. Touvron, H., Sablayrolles, A., Douze, M., Cord, M. & Jégou, H. Grafit: Learning fine-grained image representations with coarse labels. in 854–864 (IEEE Computer Society, 2021). doi:10.1109/ICCV48922.2021.00091.

22. Makeig, S., Bell, A., Jung, T.-P. & Sejnowski, T. J. Independent Component Analysis of Electroencephalographic Data. in Advances in Neural Information Processing Systems vol. 8 (MIT Press, 1995).

23. Pfurtscheller, G., Guger, C. & Ramoser, H. EEG-based brain-computer interface using subject-specific spatial filters. in Engineering Applications of Bio-Inspired Artificial Neural Networks (eds. Mira, J. & Sánchez-Andrés, J. V.) 248–254 (Springer, 1999). doi:10.1007/BFb0100491.

24. Geisler, C. et al. Temporal delays among place cells determine the frequency of population theta oscillations in the hippocampus. Proc. Natl. Acad. Sci. 107, 7957–7962 (2010).

25. Geisler, C., Robbe, D., Zugaro, M., Sirota, A. & Buzsáki, G. Hippocampal place cell assemblies are speedcontrolled oscillators. Proc. Natl. Acad. Sci. U. S. A. 104, 8149–8154 (2007).

26. Einevoll, G. T., Kayser, C., Logothetis, N. K. & Panzeri, S. Modelling and analysis of local field potentials for studying the function of cortical circuits. Nat. Rev. Neurosci. 14, 770–785 (2013).

27. Buzsáki, G. Theta Oscillations in the Hippocampus. Neuron 33, 325–340 (2002).

28. Goutagny, R., Jackson, J. & Williams, S. Self-generated theta oscillations in the hippocampus. Nat. Neurosci. 12, 1491–1493 (2009).

29. Zhang, H., Rich, P. D., Lee, A. K. & Sharpee, T. O. Hippocampal spatial representations exhibit a hyperbolic geometry that expands with experience. Nat. Neurosci. 26, 131–139 (2023).

30. Novey, M. & Adali, T. Adaptable Nonlinearity for Complex Maximization of Nongaussianity and a Fixed-Point Algorithm. in 2006 16th IEEE Signal Processing Society Workshop on Machine Learning for Signal Processing 79–84 (2006). doi:10.1109/MLSP.2006.275526.

31. Jung, T. P. et al. Removing electroencephalographic artifacts by blind source separation. Psychophysiology 37, 163–178 (2000).

32. Grosse-Wentrup, M. & Buss, M. Multiclass common spatial patterns and information theoretic feature extraction. IEEE Trans. Biomed. Eng. 55, 1991–2000 (2008).

33. Lawhern, V. J. et al. EEGNet: a compact convolutional neural network for EEG-based brain–computer interfaces. J. Neural Eng. 15, 056013 (2018).

34. Schirrmeister, R. T. et al. Deep learning with convolutional neural networks for EEG decoding and visualization. Hum. Brain Mapp. 38, 5391–5420 (2017).

35. Petersen, P. C. & Buzsáki, G. Cooling of Medial Septum Reveals Theta Phase Lag Coordination of Hippocampal Cell Assemblies. Neuron 107, 731-744.e3 (2020).

36. Colgin, L. L. et al. Frequency of gamma oscillations routes flow of information in the hippocampus. Nature 462, 353–357 (2009).

37. Hasselmo, M. E. & Stern, C. E. Theta rhythm and the encoding and retrieval of space and time. NeuroImage 85, 656–666 (2014).

38. Lustig, B. R. CONTEXT DEPENDENT ACTIVITY IN RAT HIPPOCAMPUS DURING PERFORMANCE OF A THREE ARM DELAYED SEQUENCE TASK. (2018).

39. Hazan, L., Zugaro, M. & Buzsáki, G. Klusters, NeuroScope, NDManager: a free software suite for neurophysiological data processing and visualization. J. Neurosci. Methods 155, 207–216 (2006).

40. Fan, R.-E., Chang, K.-W., Hsieh, C.-J., Wang, X.-R. & Lin, C.-J. LIBLINEAR: A library for large linear classification. J. Mach. Learn. Res. 9, 1871–1874 (2008).

41. Qin, Z., Kim, D. & Gedeon, T. Rethinking Softmax with Cross-Entropy: Neural Network Classifier as Mutual Information Estimator. Preprint at 10.48550/arXiv.1911.10688 (2020).

42. Blankertz, B., Tomioka, R., Lemm, S., Kawanabe, M. & Muller, K. Optimizing Spatial filters for Robust EEG Single-Trial Analysis. IEEE Signal Process. Mag. 25, 41–56 (2008).

43. Mitra, P. P. & Bokil, H. Observed Brain Dynamics. (Oxford University Press, 2007).

44. Markus, E. J. et al. Interactions between location and task affect the spatial and directional firing of hippocampal neurons. J. Neurosci. 15, 7079–7094 (1995).

